# A Novel *Caenorhabditis elegans* Gene Network Uncovers Mechanisms of Mitochondrial Maintenance

**DOI:** 10.1101/2022.04.06.487352

**Authors:** Armando Moreno, Allison Taffet, Elissa Tjahjono, Natalia V. Kirienko

**Author notes:** These authors contributed equally (Armando Moreno, Allison Taffet). Corresponding Author:, Mailing Address: 6100 Main St, MS-140, Houston, TX 77005.

## Abstract

Mitochondria play key roles in cellular health and metabolism and are a critical determinant of the activation of multiple cell death processes. Although several pathways for regulating and re-establishing mitochondrial homeostasis have been identified within the past twenty years, large gaps remain in our understanding of how cells keep mitochondria healthy.

To address this limitation, have developed a network of genes that underlie mitochondrial health. We began by compiling a list of frequently mutated genes using publicly available data from multiple human cancer cell lines. RNAi was used to disrupt orthologous genes in the model organism *Caenorhabditis elegans* in a series of assays to evaluate these genes’ ability to support mitochondrial health, as evidenced by precocious activation of mitochondrial autophagy and sensitivity to acute mitochondrial damage. Iterative screening of ^~^1000 genes yielded a network of 139 genes showing significant connectivity.

Functional validation of a panel of genes from the network indicated that disruption of each gene triggered at least one phenotype consistent with mitochondrial dysfunction, including increased fragmentation of the mitochondrial network, abnormal steady-state levels of ATP, NADH, or ROS, and altered oxygen consumption. Importantly, RNAi-mediated knockdown of these genes often exacerbated α-synuclein aggregation in a *C. elegans* model of Parkinson’s disease, indicating significant changes to cellular health. Additionally, human orthologs of the final mitochondrial health gene network showed enrichment for roles in a number of human disorders identified in the OMIM database. This gene network provides a foundation for identifying new mechanisms that support mitochondrial and cellular homeostasis.

## INTRODUCTION

As organisms grow and develop, their cells undergo repeated rounds of division that each require the duplication of prodigious amounts of genomic DNA. During this replication process, spontaneous mutations occur across the genome due to copying errors by DNA polymerases. Accumulation of these mutations increasingly burdens protein functions and places ever greater strain on cellular quality control systems that begin to falter as biological age increases. Failure of these quality control mechanisms, especially those involved in DNA replication and repair, often results in neoplasias that may develop into cancer (Kraemer *et al*. 1984; Koorstra *et al*. 2009; Torgovnick and Schumacher 2015). On this basis, cancerous tissues may represent a relatively untapped resource for the identification of previously underappreciated programs for the maintenance of cellular health.

Mitochondria are a critical component for cellular health, and a key determinant of the ability of apoptosis to limit the proliferation of cancer cells. Most commonly known as the “powerhouse of the cell”, mitochondria are much more than just a site for energy production; they participate in a myriad of functions, including several cell death pathways (Wang and Youle 2009; Rudel *et al*. 2010; Marquez-Jurado *et al*. 2018), homeostatic regulation of calcium and iron (Zhang *et al*. 2019), coordinating the production of reactive oxygen species (ROS) (Li *et al*. 2013; Zorov *et al*. 2014; Dan Dunn *et al*. 2015), and providing essential intermediates and regulation of cholesterol biosynthesis (Pignataro *et al*. 1983). Several critical pathways for maintaining mitochondrial homeostasis, including surveillance, repair, and recycling, have been identified within the past few decades, and often transcend this function to support general cellular homeostasis as well (Yuan *et al*. 2013; Jovaisaite *et al*. 2014; Tjahjono and Kirienko 2017). For example, recent work from our lab has shown that a mitochondrial surveillance pathway called the ESRE network is activated by an increase in ROS is crucial for both mitochondrial and cellular homeostasis (Tjahjono *et al*. 2020). Despite these discoveries leading to a greater understanding of mitochondrial maintenance, gaps remain in our understanding of how cellular aging affects mitochondrial function.

With so many genes being involved in mitochondrial maintenance, an unbiased screen for genes exhibiting an effect on mitochondrial response to stress is a valuable approach to better understand the mechanistic relationships between mitochondrial homeostasis and cellular health. The model organism *Caenorhabditis elegans* provides an invaluable tool to conduct high-throughput genetic studies. More specifically, *C. elegans* is very useful in reverse genetic screens through the ease of creating gene knockdowns by the feeding of RNAi (Conte *et al*. 2015). Not only does this model organism lend itself to reverse genetic studies, but high-throughput protocols have been well established over the years (O’Reilly *et al*. 2014; Anderson *et al*. 2018; Sohrabi *et al*. 2021). This high-throughput system, in conjunction with many available *C. elegans* reporter lines, including those related to mitochondrial health, including PINK-1::GFP translational fusion that allows to monitor activation of mitophagy (Tjahjono *et al*. 2020), is highly adaptable to screen for genes related to mitochondrial function.

In this study, we used the NCI-60 cancer cell database to conduct a high-throughput genetic screen of frequently mutated genes mutated in cancer. As mutagenesis process is not entirely random, and some genes mutate with higher frequency than others, this allowed us to identify a set of frequently-mutated genes, regardless of their tissue origin. This set included both cancer driver and passenger mutations. We evaluated whether disrupting these genes affected sensitivity to mitochondrial damage. We developed a gene network comprised of 139 genes that have a significant number of connections that spans a wide variety of biological functions and pathways. Subsequent validation of a subset of these genes demonstrated that their knockdown results in the aberrant mitochondrial morphology and / or function. In addition, negative phenotypic outcomes, including slower pumping, smaller brood size, or increased protein aggregation were observed.

## MATERIALS AND METHODS

### Media and Strains

Standard conditions were used in the maintenance of *C. elegans*. Worms were propagated on *Escherichia coli* OP50-seeded NGM plates at 20 °C, synchronized by hypochlorite bleaching, and hatched overnight at room temperature. Synchronized worms were maintained in S Basal at 15 °C prior to experimental use. The following *C. elegans* strains were used: N2 Bristol (wild-type), SS104 [*glp-4(bn2)*], NVK90 [*pink-1(tm1779); houIs001*], SJ4103 [*zcIs14/Pmyo-3*::GFPmt)], PE255 [*sur-5p*::luciferase::GFP + *rol-6(su1006)*], ALF86 (*Pmyo-3*::Peredox::*unc-119*)and NL5901 [*pkIs2386* (P*unc-54*::α-synuclein::YFP *unc-119*(+))]. For all RNAi experiments, worms were reared on NGM/Carbenicillin(25μg/ml)/IPTG(1mM) plates containing HT115(DE3)-based RNAi strains for 68 hours at 20 °C unless otherwise noted. RNAi clones all came from the Ahringer or Vidal libraries and were sequenced for verification.

### Precocious Mitophagy Activation (Primary Screening Assay)

^~^ 200 L1 stage NVK90 worms were dropped onto 6 cm NGM/Carb/IPTG, RNAi seeded plates and incubated at 25 °C for 48 hours. Each 6 cm plate was washed with S Basal and 200 μl of worms were transferred into two wells of a 96 well plate. The worms from the 96 well plate were sorted into a 384 well plate using a COPAS FlowPilot and Large Particle (LP) Sampler, with 20 worms per well. 7 mM sodium selenite and HT115 bacteria (food source, final OD600=0.1) were added to the 384-well plates. Brightfield and GFP images were taken every 3 hours for 30 hours using the Cytation 5 Cell Imaging Multi-Mode Reader (BioTek Instruments). GFP quantification was performed by a Cell Profiler program (https://cellprofiler.org/). Gene knockdowns were counted as hits if the GFP level was 1.5 times that of vector control. At least three biological replicates were performed.

### Sensitivity to Acute and Chronic Mitochondrial Damage (Secondary Screening Assay)

For measurement of sensitivity to acute mitochondrial damage, L1 stage SS104 worms were dropped onto 6 cm NGM/Carb/IPTG, RNAi-seeded plates and incubated overnight at 20 °C then shifted to 25 °C for 48 h. Worms were placed into 384-well plates as described above. 0.357 μM Sytox Orange stain (Invitrogen) was added to each well and incubated for 5 hours. Then, 50 μM of an uncoupler CCCP (Sigma) was added to each well. Brightfield and RFP-channel images were taken every 30 minutes for 12 hours using the Cytation 5 Cell Imaging Multi-Mode Reader (BioTek Instruments). Fluorescence quantifications and the counting of fraction of dead worms were performed by a Cell Profiler program. Gene knockdowns were counted as hits if the number of dead worms was 1.5 times that of the empty vector control.

For measurement of sensitivity to chronic mitochondrial damage, L1 stage SS104 worms were dropped onto 6 cm NGM/Carb/IPTG, RNAi-seeded plates and incubated at 25 °C for 72 hours. Worms were picked onto 35 mm OP50-seeded NGM plates supplemented with 100 μM of iron chelator 1,10-phenanthroline (Sigma). Four days later, worms were scored and counted as dead if they did not respond to touch. Three biological replicates were performed, and each biological replicate consisted of three technical replicates, each of 60 worms. RNAi strains were compared to the empty vector control and genes that showed a significant increase in death compared to the empty vector control were considered hits. Student’s t-test was used to calculate statistical significance.

### Longevity Assay (Tertiary Screening Assay)

During first iteration of screening (seed set determination), for measurement of longevity SS104 worms were grown to a young adult stage on RNAi or vector control plates as described above. Then they were placed on OP50 NGM plates and were scored every other day. Three biological replicates comprised of three technical replicates each, were performed (^~^ 150 worms/biological replicate). Log-rank test was used to determine statistical significance. For the second iteration of screening (network expansion), worms were grown and longevity plates were set as above, but survival on a single day, day 8, was scored. RNAis with the survival of 90% of vector control were considered healthy (i.e. if 85% of C. elegans on vector RNAi were alive, RNAi with survival > 76.5% will be considered healthy).

### Fluorescence Microscopy

To visualize the mitochondrial network morphology, L1 stage SJ4103 worms were grown on 6 cm NGM/Carb/IPTG, RNAi-seeded plates and incubated at 20 °C for 72 h. The worms were then washed in S Basal, immobilized with levamisole (Sigma) and pipetted onto agar pads. At least 30 worms were used per condition per biological replicate, and three replicates were completed per condition. Images were acquired using a Zeiss Axio Imager M2 upright microscope with a Zeiss AxioCam 506 Mono camera and Zeiss Zen2Pro software. Statistical significance was determined by chi-squared test.

To visualize alpha-synuclein aggregation L1 stage NL5901 worms were grown on 6 cm NGM/Carb/IPTG, RNAi seeded plates and incubated at 20 °C for 72 h. The worms were then picked onto 6 cm OP50-seeded NGM plates and incubated at 20 °C for 48 h. Worms were then washed in S Basal, immobilized with levamisole (Sigma) and pipetted onto agar pads. At least 25 worms were used per condition per biological replicate, and six replicates were completed per condition. Worms were visualized using a Zeiss Axio Imager M2 upright microscope with a Zeiss AxioCam 506 Mono camera and Zeiss Zen2Pro software and aggregates were manually counted.

### ATP Measurements

To measure steady state ATP levels, L1 stage PE255 worms were grown on 6 cm NGM/Carb/IPTG, RNAi seeded plates and incubated at 20 °C for 72 h. The worms were then washed in S Basal and transferred to a 96-well plate (50 worms per well, 4 wells per RNAi condition). 0.3 mM luciferin buffer was added, and the plate was loaded into the Cytation 5 Cell Imaging Multi-Mode Reader (BioTek Instruments), where luminescence and GFP fluorescence values were read. GFP was used for protein normalization purposes.

### NADH/NAD+ Ratio Measurements

To measure NADH/NAD+ ratio, L1 stage ALF86 worms were grown on 6 cm NGM/Carb/IPTG, RNAi seeded plates and incubated at 20 °C for 72 h. The worms were then washed in S Basal and transferred to a 96-well plate (50 worms per well, 4 wells per RNAi condition). The plate was then loaded into the Cytation 5 Cell Imaging Multi-Mode Reader (BioTek Instruments) and fluorescence in GFP and RFP channels was obtained. The ratio was calculated based on the GFP and RFP fluorescence ratios.

### Flow Vermimetry

L1 stage N2 worms were grown on 6 cm NGM/Carb/IPTG, RNAi seeded plates and incubated at 20 °C for 72 h. Worms were then washed with S Basal, transferred to 96-well plates, and stained with 10 μM of 10-nonyl acridine orange bromide (Invitrogen) for mitochondrial mass or 4.5 μM MitoTracker Red CMXRos (Invitrogen) for membrane potential, in S Basal and compared to no-stain controls. Fluorescence was measured with a COPAS FlowPilot and normalized to time of flight to account for worm size. Three biological replicates were performed, and each consisted of two technical replicates, with 200 worms each. Worm size data was recorded by the COPAS FlowPilot through TOF and normalized to the empty vector control.

To measure ROS levels, L1 stage SS104 worms were gown on 6 cm NGM/Carb/IPTG, RNAi seeded plates and incubated at 20 °C for 24 h then shifted to 25 °C for 48 h. Worms were then washed with S Basal, transferred to 96-well plates, and stained with 3 μM of dihydroethidium (DHE, Thermo Fisher). The plates were incubated in the dark at room temperature for 1 h and then were washed with S Basal to remove residual dye. Fluorescence was measured with a COPAS FlowPilot and normalized to time of flight to account for worm size. Three biological replicates were performed, and each consisted of three technical replicates, with 400 worms each.

### Oxygen Consumption

10,000 worms were grown on 10 cm NGM/Carb/IPTG, RNAi seeded plates and incubated at 20 °C for 72 h. Worms were washed in S Basal and then counted and sorted using the COPAS FlowPilot. A max of 10,000 worms were used, with an average of 8,000 worms per replicate, and the exact worm number was recorded. 30 minutes were allowed before measurement for digestion of bacteria. Oxygen consumption was measured at 20 °C with a YSI Model 5300 Biological Oxygen Monitor and a YSI 5301 Clark-type oxygen electrode (Yellow Springs Instrument), and readings were continuously recorded for about 5-15 minutes, depending on rate. Oxygen consumption rate was measured from the slope of the straight piece of the recorded graph and normalized to worm number. 5-7 replicates were completed for each condition.

### Statistical Analysis

One-way ANOVA analysis was used to calculate significance unless otherwise stated. Dunnett’s post hoc analysis was used to calculate *p*-values between each condition and the control group. On the graphs statistical significance is represented as follows: \**p*Ͱ< 0.05, \*\**p*Ͱ<Ͱ0.01, and \*\*\**p*Ͱ<Ͱ0.001.

## RESULTS

### Identification of a gene network associated with increased mitochondrial sensitivity

The NCI-60 cancer database is comprised of 60 different tumor cell lines spanning a wide variety of tumor type and full genomic sequence data. Using data within the NCI-60 database, we used the Catalogue of Somatic Mutations in Cancer (COSMIC) and the Cancer Cell Line Encyclopedia (CCLE) databases to identify 15,856 genes with mutations in at least one cell line. To improve robustness, any gene that was not mutated in at least 10 cell lines originating from at least three different tissues was removed, leaving us with 385 genes. The list was also supplemented with genes that were mutated in at least three of the eight NCI-60 cell lines that were most sensitive to mitochondrial damage (LE:SR, LE:CCRF-CEM, ME:LOX IMVI, LE:MOLT-4, CNS:U251, BR:MCF7, LE:HL-60(TB), LC:NCI-H460) (Panina *et al*. 2019). This added a total of 251 genes to the list and was expected to enrich the list for genes that played a role in mitochondrial sensitivity to damage. Finally, 20 genes known to be implicated in either tumor suppression or promotion based on the literature were manually added. Bioinformatic tools, such as Ortholist and Ensembl (Hubbard *et al*. 2002; Kim *et al*. 2018), were used to identify putative *C. elegans* orthologs for each of the 656 human genes, resulting in 585 genes that were tested. A testing pipeline was developed that would allow identification of genes that play a role in mitochondrial health **(Figure 1A)**.

**Figure 1.**
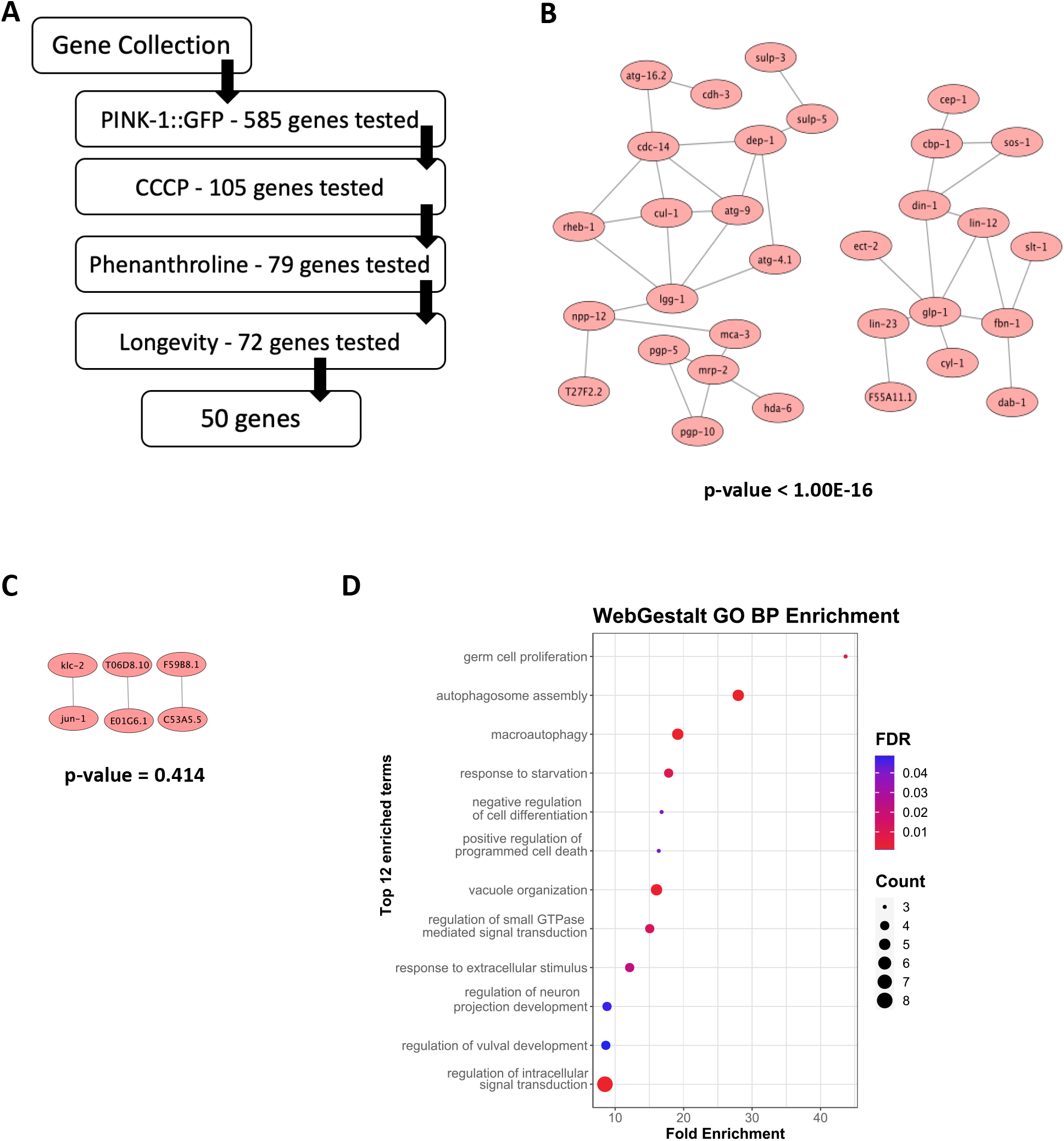
Identification of a gene network underlying mitochondrial health. **(A)** Flowchart of testing process for genes identified from the NCI-60 hit list. The flowchart indicates the number of genes tested at each stage. **(B)** Visualization of the initial 50 gene ‘seed set’, showing 38 connections between the genes. **(C)** Connectivity of one representative set (of 100 sets) of 50 randomly selected genes from the *C. elegans* genome, showing a total of 3 connections. *p*-values **(B, C)** were calculated by the WormNetV3 database and include a Bonferroni correction. **(D)** Bioinformatic analysis of the 50 gene set showed enrichment for cell stemness (e.g., germ cell proliferation and negative regulation of differentiation) and autophagic flux (autophagosome assembly, macroautophagy, response to starvation, etc.)

RNAi was used to knock down each of the genes in a *C. elegans* reporter strain carrying a P*pink*-1::PINK-1::GFP translational reporter. Synchronized L1 worms were dropped onto *E. coli* expressing dsRNA of the gene of interest and incubated for 48 hours in 25 °C. Early L4 stage worms were then exposed to sodium selenite (a known inducer of mitophagy). Brightfield and fluorescence images were collected every 2 h from 10 to 24 h after exposure to selenite to observe PINK-1::GFP accumulation. Images were then analyzed by an unbiased, automated pipeline in CellProfiler to compare fluorescence levels in vector controls to RNAi mutants. Specifically, PINK-1::GFP accumulation was compared once 10-20% of vector control worms exhibited clear signs of PINK-1::GFP accumulation. This provided some range for increased PINK-1::GFP accumulation, indicating that the gene disruption caused precocious mitophagic activation. Knockdowns that exhibited at least 1.5-fold increase in the number of worms expressing PINK-1::GFP were considered hits in the primary assay **(Supplemental Figure S1A)**.

105 genes (16% of the original set) passed this primary cutoff and were moved to a secondary screen, where RNAi was used to test whether knockdown increased sensitivity to acute or chronic mitochondrial stress. For these tests, synchronized L1, temperature-sensitive *glp-4(bn2ts)* worms were reared on RNAi for each of the 105 genes. The *glp-4(bn2)* allele results in a temperature-dependent sterility that simplifies lifespan and survival assays. For acute mitochondrial damage, young adult worms were sorted into 384-well plates and exposed to the mitochondrial uncoupler carbonyl cyanide *m*-chlorophenyl hydrazone (CCCP) which causes death by depolarizing the mitochondrial membrane (Chaudhari *et al*. 2008). Starting 6 h after exposure to CCCP, survival was assayed every 30 min for 6 h using a vital dye Sytox Orange. Genes whose disruption increased death by at least 1.5-fold compared to the vector control were considered hits **(Supplemental Figure S1B)**.

79 genes (75% of the genes tested in the secondary screen) met this criterion and were tested in an orthogonal screen for increased sensitivity to chronic mitochondrial damage. Young adult *glp-4(bn2)* worms were placed on NGM plates containing 100 μM 1,10-phenanthroline. Phenanthroline is a small molecule iron chelator that damages mitochondria and kills *C. elegans* (de Avellar *et al*. 2004). Survival was scored daily until vector controls reached at least 90% death. Log-rank tests were used to identify genes whose disruption significantly reduced host survival. 72 of the 79 genes tested (^~^90%) had this effect.

To eliminate the possibility that increased sensitivity was due to the hit genes being required for survival under normal conditions, lifespan assays were performed with RNAi of the 79 genes in *glp-4(bn2)* worms. Genes whose disruption reduced survival below 90% of vector control were removed from the list. 29 genes (37%) of the genes were removed for this reason.

These experiments yielded an initial seed list of 50 genes (see **Supplemental Table S1** for the gene list). Gene interaction assessment tools, such as String and WormNet (a network-assisted predictive tool that mines dozens of connection types, including *in vivo* and *in vitro* protein-protein interactions, genetic interactions, interactions between orthologs, co-citations, etc) identified 38 connections amongst these 50 genes **(Figure 1B)**. There was significantly greater connectivity (*p-value*<1.00E-16) than was observed, on average for 100 networks of 50 randomly-chosen genes (*p-value*=0.414) **(Figure 1C)**.

This left an initial seed list of 50 genes associated with membrane transport, stemness (e.g., germ cell proliferation and negative regulation of differentiation), and autophagic flux (autophagosome assembly, macroautophagy, response to starvation, etc.), indicating that our gene list spans a variety of biological functions **(Figure 1D)**.

### Expansion of the network identifies genes that impact mitochondrial health

This seed set was useful, as all of its genes demonstrated mitochondria-related phenotypes. However, it was fairly small. To expand this list, WormNet was used to identify additional genes that were significantly (*p*-value<0.05) associated with our seed set. This yielded ^~^400 additional candidate genes **(Supplemental Table S2)**. After removing genes whose mutation is lethal, 381 genes remained. Similar to seed set, RNAi was used to test each of these genes for PINK-1::GFP accumulation, increased sensitivity to an acute mitochondrial poison, and lifespan analyses. Analysis of the initial screening set showed a very strong correlation between acute and chronic mitochondrial toxicity (90% of hits in acute sensitivity assay were also hits in chronic sensitivity assay), so the latter assay was dropped for further analysis.

89 of the 381 expansion genes demonstrated increased mitochondrial sensitivity without general malaise, giving an expanded network of 139 genes **(Figure 2A-2C)**. Using WebGestalt, we discovered a 6-fold enrichment in genes associated with cell division **(Figure 2D)**. This enrichment is likely due to the use of cancer cells for the initial identification, but our results also suggest a connection between cell cycle genes and mitochondrial dysfunction. Genes associated with the ErbB (e.g., *mkk-4, plc-3, nck-1, sem-5, ced-2*, and *sos-1*) and Wnt (e.g., *kin-3, kin-10, cul-1, lin-23, pop-1, cbp-1, bar-1, cyd-1*, and *rho-1*) signaling pathways were also enriched. Both of these pathways are associated with cancer (Yarden and Sliwkowski 2001; Zhan *et al*. 2017). While these pathways have been studied extensively, there is still little known about how they contribute to mitochondrial health.

**Figure 2.**
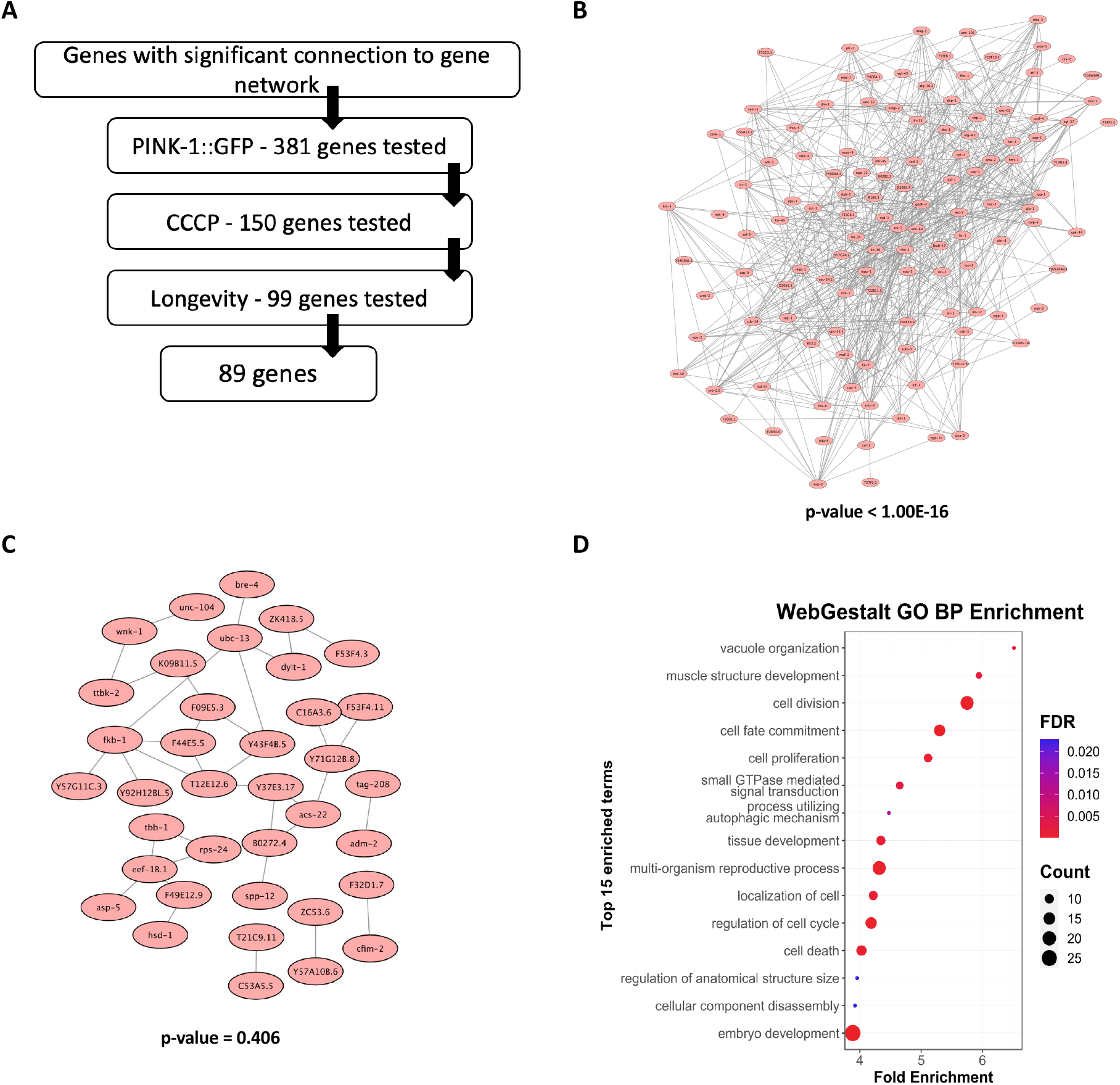
Refinement of the mitochondrial health gene network. **(A)** Flowchart for testing the 381 new genes identified by their connection to the seed set. **(B)** Connectivity of the expanded gene network 139 genes. **(C)** Connectivity of one representative set (of 100 sets) of 140 randomly selected genes from the *C. elegans* genome, showing a total of ^~^40 connections. *p*-values **(B, C)** were calculated by the WormNetV3 database and include a Bonferroni correction. **(D)** Bioinformatic analysis of the 139 gene set showed enrichment for cell proliferation and development as well as signal transduction.

### Disruption of network members impairs mitochondrial function

To examine the consequences of gene knockdown on mitochondrial function, a panel of 8 genes with diverse functions and cellular locations (*ogdh-1, kin-10, ego-2, bar-1, pps-1, exos-9, cdc-37*, and *cul-1*) were selected for further study. These genes span a wide variety of biological functions; some are known to have involvement with mitochondrial function while others do not **(Table 1)**.

**Table 1.** A description of panel of 8 network members. Ten genes were chosen to represent the gene network as a whole. These included genes with both known and presumed (on the basis of this network) to have functions in mitochondrial health maintenance.

| Gene in *C. elegans* | Protein | Human Orthologue | Direct involvement with mitochondrial function? | Biological Function |
| --- | --- | --- | --- | --- |
| *ogdh-1* | 2-oxoglutarate dehydrogenase | OGDH | Yes | Involved in the conversion of 2-oxoglutarate to succinyl-CoA |
| *kin-10* | Casein kinase II subunit beta | CSNK2B | No | Modulates TRPP function (Hu *et al.* 2006)<br>Involved in Wnt signaling (DE GROOT *et al.* 2014) |
| *ego-2* | BRO1 domain-containing protein | PTPN23 | No | Promotes Notch signaling (LIU AND MAINE 2007) |
| *bar-1* | Beta-catenin/armadillo-related protein 1 | CTNNB1 | No | Regulates Hox gene *lin-39* expression through Wnt signaling (EISENMANN *et al.* 1998) |
| *pps-1* | 3'-phosphoadenosine-5'-phosphosulfate synthase | PAPSS1 | Yes | Involved in sulfate metabolism (DEJIMA *et al.* 2006) |
| *exos-9* | Exosome complex component RRP45 | EXOSC9 | No | Not well studied but exhibits exonuclease activity |
| *cdc-37* | Probable Hsp90 co-chaperone cdc37 | CDC37 | No* | Cofactor for many kinases and is involved in ATPase activity (ECKL *et al.* 2013) |
| *cul-1* | Cullin-1 | CUL1 | No | Involved in protein degradation (GHAZI *et al.* 2007) |

\*While CDC37 direct interaction with mitochondria has not been observed, its probable co-chaperone HSP90 has.

As a first step, we used a *C. elegans* reporter strain that carries a mitochondrially-targeted GFP expressed in body wall muscle cells (referred to herein as mtGFP). This strain allows for ready visualization of the mitochondrial network in these cells. Normally in *C. elegans* these mitochondria are hyperfused and form a network comprised of long, branched tubules. Mitochondrial damage causes this network to fragment. For this test, synchronized L1 mtGFP worms were reared on RNAi targeting each of the 8 genes. Once the worms reached the L4 stage, fluorescence microscopy was used to visualize and quantify mitochondrial fragmentation. RNAi for each of the genes resulted in significant mitochondrial fragmentation (*p-value*<0.001), supporting the conclusion that they are associated with mitochondrial maintenance **(Figure 3)**.

**Figure 3.**
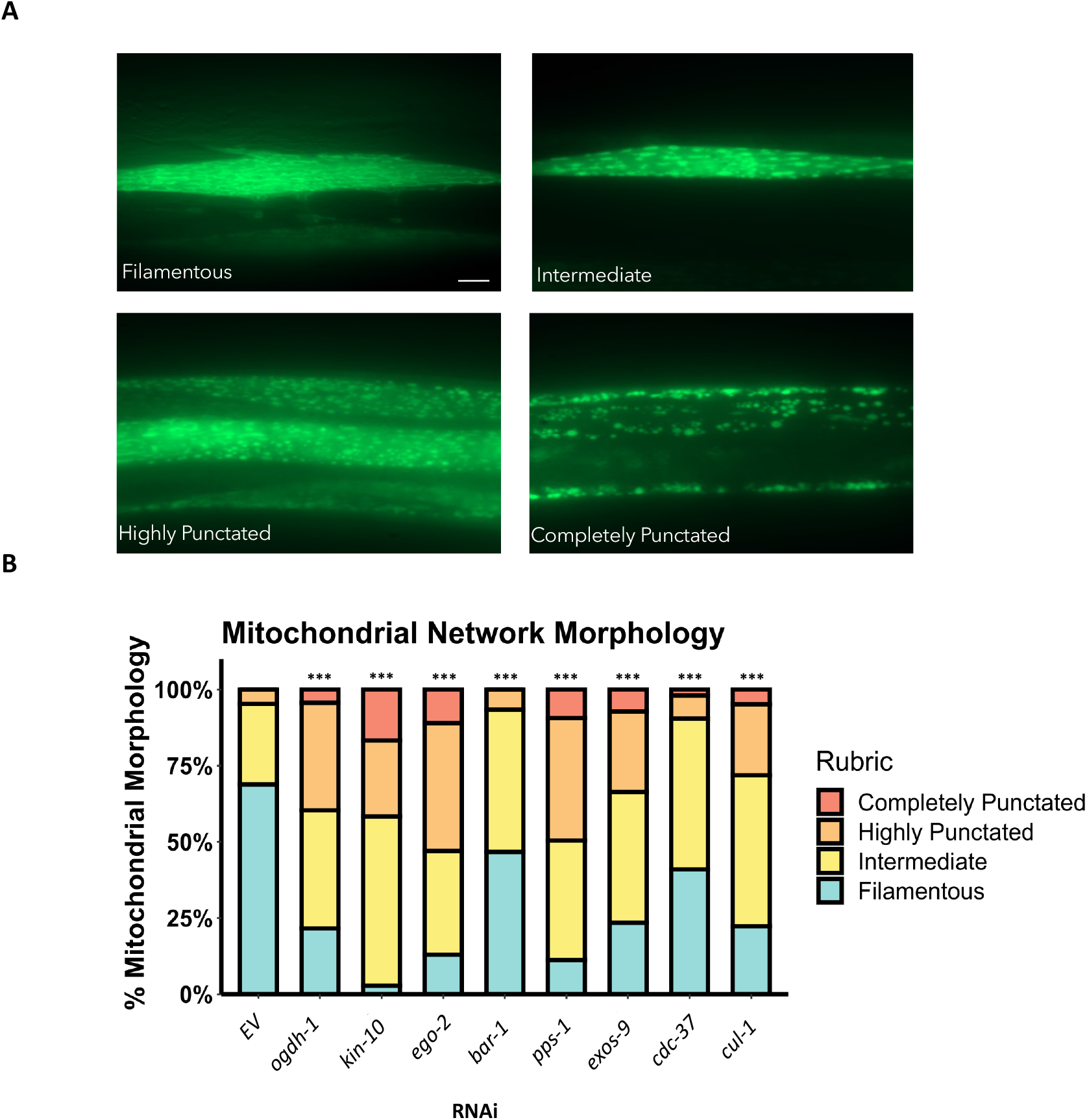
Genes in the mitochondrial health gene network are support mitochondrial network connectivity. **(A)** Fluorescence micrographs of mitochondrially-targeted GFP (mtGFP) expressed in body wall muscle cells showing four different qualitative states. Shown are filamentous (top left), intermediate fragmentation (upper right), highly punctate (bottom left) and completely punctate (bottom right). **(B)** Quantification of fragmentation state of worms expressing mtGFP in body wall muscles after RNAi targeting selected genes. ^~^90 worms were examined for each genotype. All genes showed an increase in mitochondrial fragmentation after RNAi. Statistical significance was determined by χ^2^ analysis, ***p<0.001

Next, a set of reporter strains and fluorescent dyes was used to assess mitochondrial function after gene knockdown. Worms carrying an ATP-dependent luciferase were used to assess steady state levels of ATP. In parallel, we used the Peredox reporter, which fluoresces due to an NADH-dependent conformational change and can be used to obtain a semi-quantitative readout of cellular reductive potential. Synchronized L1 worms carrying either reporter were spotted onto RNAi of the eight genes in the panel and reared to the L4 stage and then ATP and NADH levels were determined, respectively. For ATP, only *ogdh-1* showed reductions, while *kin-10(RNAi), ego-2(RNAi)*, and *pps-1(RNAi)* actually showed more ATP than vector controls **(Figure 4A)**. *ogdh-1(RNAi), ego-2(RNAi)* and *pps-1(RNAi)* disrupted redox status, as demonstrated by the increase in fluorescence of the Peredox reporter **(Figure 4B)**.

**Figure 4.**
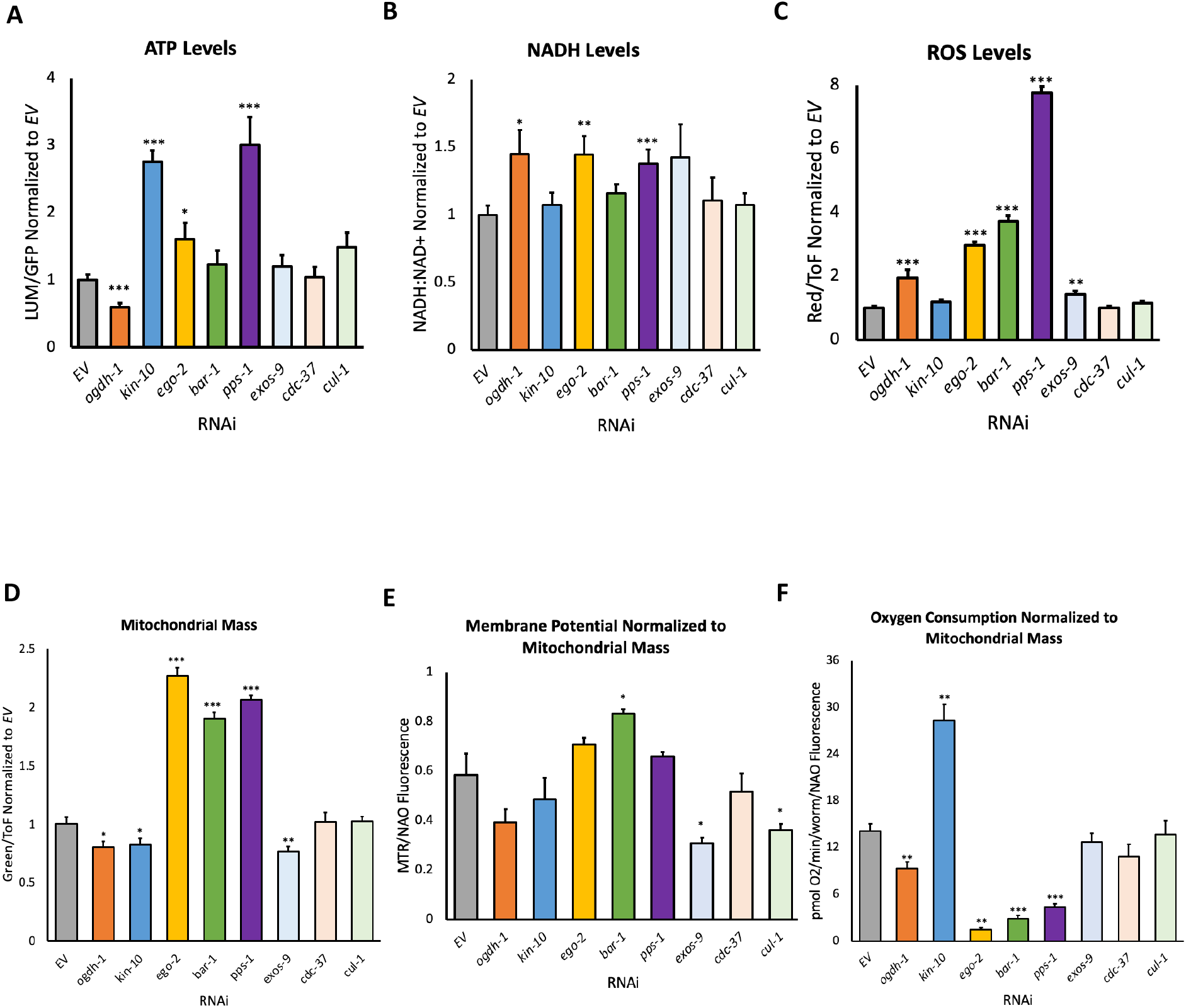
Genes in the mitochondrial health gene network support mitochondrial health. **(A)** Bar graph of steady-state ATP levels in young adult worms reared on the indicated RNAi constructs. ATP levels were measured by normalizing luciferase luminescence to GFP expression in a reporter strain carrying both reporters. **(B)** Bar graph of steady-state ratio of NADH-NAD+ using a conformation-dependent fluorometric reporter. Young adult worms carrying the reporter were reared on the indicated RNAi constructs. **(C)** Bar graph of steady-state reactive oxygen species, as measured by conversion of DHE to a fluorescent state in young adult worms reared on RNAi targeting the indicated gene. Fluorescence was measured using flow vermimetry. **(D)** Bar graph of mitochondrial mass, as measured by mitochondrial staining with 10-*N*-nonyl acridine orange (NAO) in young adult worms reared on RNAi targeting the indicated gene. Fluorescence was measured using flow vermimetry. **(E)** Bar graph of mitochondrial membrane potential, as measured by the dye MitoTracker Red CMXRos, normalized to mitochondrial mass, as measured by NAO, in young adult worms reared on RNAi targeting the indicated gene. Fluorescence was measured using flow vermimetry. **(F)** Bar graph of basal oxygen consumption in young adult worms reared on RNAi targeting the indicated gene as measured by a Clark-electrode. Statistical significance in all panels was calculated using one-way ANOVA analysis followed by a Dunnett’s post-hoc test. *p<0.05, **p<0.01, ***p<0.001

Steady-state ROS levels were measured in young adult worms that were spotted on RNAi as synchronized L1 larvae. Upon reaching the young adult stage, worms were stained with the ROS-reactive dye dihydroethidium (DHE), and fluorescence (normalized to worm size) was measured using flow vermimetry in a COPAS flow pilot (Tjahjono *et al*. 2021). Five knockdowns: *ogdh-1(RNAi), ego-2(RNAi), bar-1(RNAi), pps-1(RNAi)*, and *exos-9(RNAi)* showed increased ROS **(Figure 4C)**.

Mitochondrial mass and membrane potential were measured using the fluorescent dyes 10-nonyl acridine orange and MitoTracker Red, respectively. Mitochondrial mass was significantly increased in *ego-2(RNAi), bar-1(RNAi)* and *pps-1(RNAi)* **(Figure 4D)**. It is worth noting that *ego-2(RNAi)* and *pps-1(RNAi)* exhibit highly fragmented mitochondrial networks, often interpreted as an imbalance between mitochondrial fission and fusion. This imbalance is known to reduce mitochondrial efficiency (Wu *et al*. 2011). Increased mitochondrial mass in these cases may be an attempt by the cells to compensate for reduced mitochondrial function. Mitochondrial membrane potential was significantly affected across all non-vector RNAi conditions **(Supplemental Figure S2)**, but this effect nearly disappeared when results were normalized to mitochondrial mass **(Figure 4E)**. For most of the knockdowns tested, the observed decrease in mitochondrial membrane potential probably results from reduced mitochondrial mass.

Next, we assessed oxygen consumption using a biological oxygen monitor and a Clark-type oxygen electrode. Oxygen consumption was measured continuously for 5-15 minutes, and basal oxygen consumption rate was derived from the slope of the recorded graph and normalized to worm count and mitochondrial mass. RNAi targeting *ogdh-1, ego-2, bar-1*, and *pps-1* resulted in significant reductions in oxygen consumption. In contrast, oxygen consumption was significantly increased for *kin-10(RNAi)* **(Figure 4G)**. Summary of these phenotypes is provided in **Table 2**.

**Table 2.** Summary table of mitochondrial phenotypes.

| Gene in *C. elegans* | ATP | NADH | ROS | Mitochondrial Mass | Membrane Potential | Oxygen Consumption | Mitochondrial Fragmentation |
| --- | --- | --- | --- | --- | --- | --- | --- |
| *ogdh-1* | ▼ | ▲ | ▲ | ▼ | — | ▼ | ▲ |
| *kin-10* | ▲ | — | — | ▼ | — | ▲ | ▲ |
| *ego-2* | ▲ | ▲ | ▲ | ▲ | — | ▼ | ▲ |
| *bar-1* | — | — | ▲ | ▲ | ▲ | ▼ | ▲ |
| *pps-1* | ▲ | ▲ | ▲ | ▲ | — | ▼ | ▲ |
| *exos-9* | — | — | ▲ | ▼ | ▼ | — | ▲ |
| *cdc-37* | — | — | — | — | — | — | ▲ |
| *cul-1* | — | — | — | — | ▼ | — | ▲ |

▲ Denotes a significant increase
▼ Denotes a significant decrease
— No significant effect

### Disruption of novel mitochondrial maintenance genes affects health in C. elegans

As noted above, while RNAi of these genes did not adversely affect lifespan **(Figure 5A)**, we wanted to assess the effects of these disruptions across a wider context of the organismal phenotypes. That could provide a valuable information on the larger effect of mitochondrial dysfunction caused by the loss of these genes. Toward this end, we measured worms’ size, fecundity, and pharyngeal pumping.

**Figure 5.**
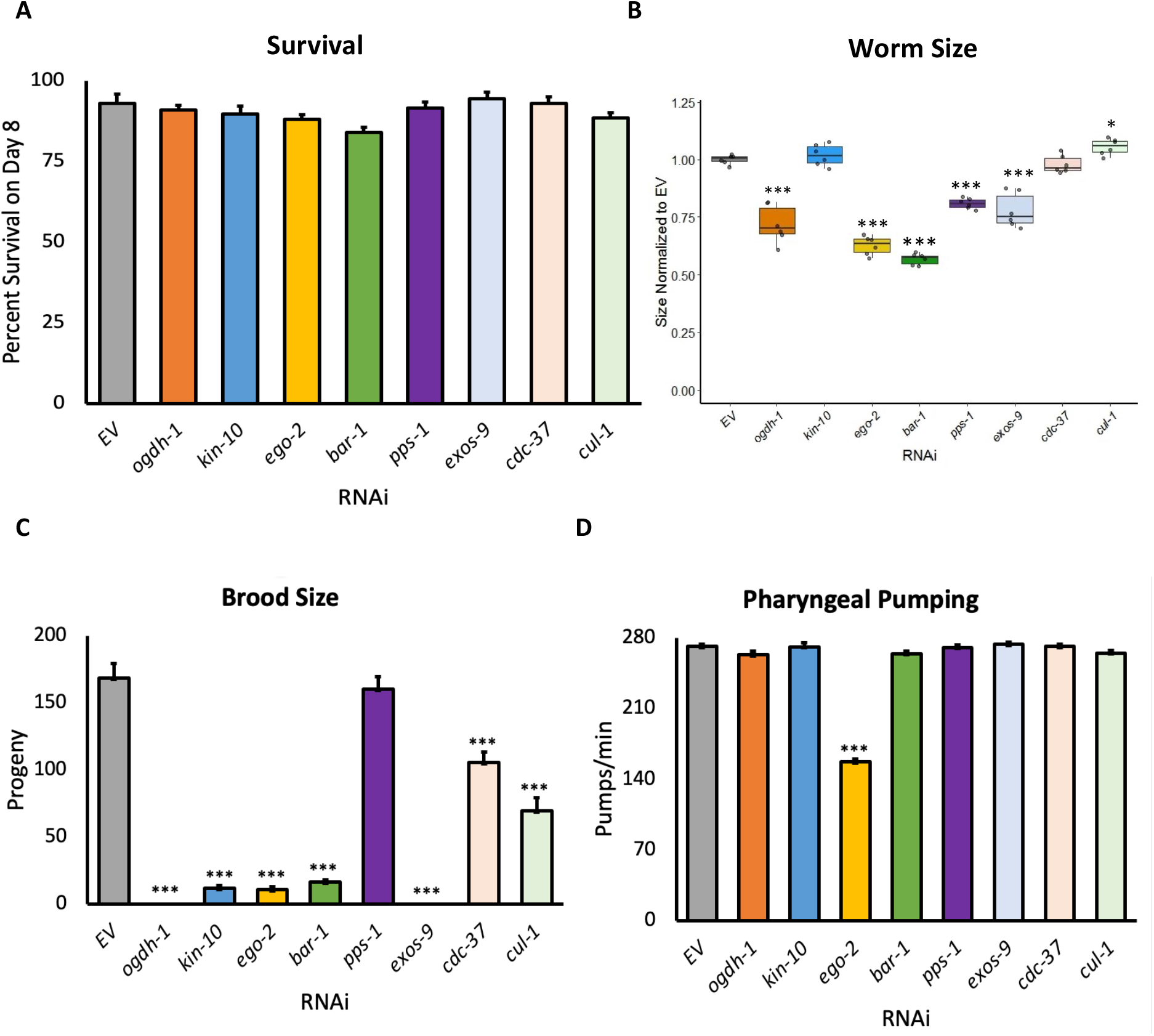
Genes in the mitochondrial health gene network support organism-level health parameters. **(A)** Bar graph of 8 d survival of worms reared on RNAi targeting the indicated gene. **(B)** Quantification of length of young adult worms reared on RNAi targeting the indicated gene. Worm length was measured using flow vermimetry. **(C)** Bar graph of worm fecundity in adult worms reared on RNAi targeting the indicated gene. **(D)** Bar graph of pharyngeal pumping rates for young adult worms reared on RNAi targeting the indicated gene. Statistical significance was calculated using one-way ANOVA analysis followed by a Dunnett’s post-hoc test. *p<0.05, **p<0.01, ***p<0.001

Size was determined by rearing synchronized L1 larvae on RNAi targeting each gene and then using the COPAS FlowPilot to measure adults using time-of-flight data. Five of the RNAi conditions reduced adult worm size, including *ogdh-1, ego-2, bar-1*, and *pps-1*, which were the most frequently affected in mitochondrial function assays **(Figure 5B)**. *exos-9(RNAi)* also had a reduced adult size. Mitochondrial damage has been shown to negatively affect fecundity as well (Zhu *et al*. 2019). To determine whether these genes have a role in this process, synchronized L1 *glp-4(bn2ts)* worms were reared on RNAi targeting the eight genes in the panel and allowed to develop to the young adult stage at permissive temperatures. Individual worms were then picked onto fresh OP50 plates and shifted to a non-permissive temperature to sterilize the offspring. Brood sizes were scored across a 5-day span, which is typically sufficient for worms to produce most of their offspring. Interestingly only *pps-1(RNAi)* retained normal fecundity **(Figure 5C)**. This result ran somewhat counter to its otherwise generally strong impact on mitochondrial phenotypes.

Finally, worms reared on each RNAi strain were observed under a dissecting microscope to score pharyngeal pumping (number of pumps per minute was manually counted). Pumping rate was only significantly decreased in *ego-2(RNAi)*, suggesting that the mitochondrial dysfunction related to disrupting these genes did not have significant effect on the muscles involved in pharyngeal pumping, thus food consumption should not be an issue from what we have seen **(Figure 5D)**.

### Genes from the network show an increase in alpha-synuclein aggregation, a feature of Parkinson’s disease

Mitochondrial dysfunction is known to be implicated in neurodegenerative diseases (NDD), and genes related to Alzheimer’s and Parkinson’s disease are also present within our network. For this reason, we tested the effects of our novel mitochondrial maintenance genes on neurodegeneration. A staple of Parkinson’s disease (PD) is the aggregation of alpha-synuclein in the form of Lewy bodies found in the brains of many patients with PD (Xu and Pu 2016). We used the NL5901 reporter strain that contains human alpha-synuclein protein fused to fluorescent reporter (YFP) expressed under control of muscle-specific promoter *Punc-54* (van Ham *et al*. 2008) to test the effects of RNAi knockdown on aggregation formation. Our results showed a significant increase in the number alpha-synuclein aggregates across most of the genes tested except for *ogdh-1(RNAi)* and *cul-1(RNAi)* **(Figure 6)**.

**Figure 6.**
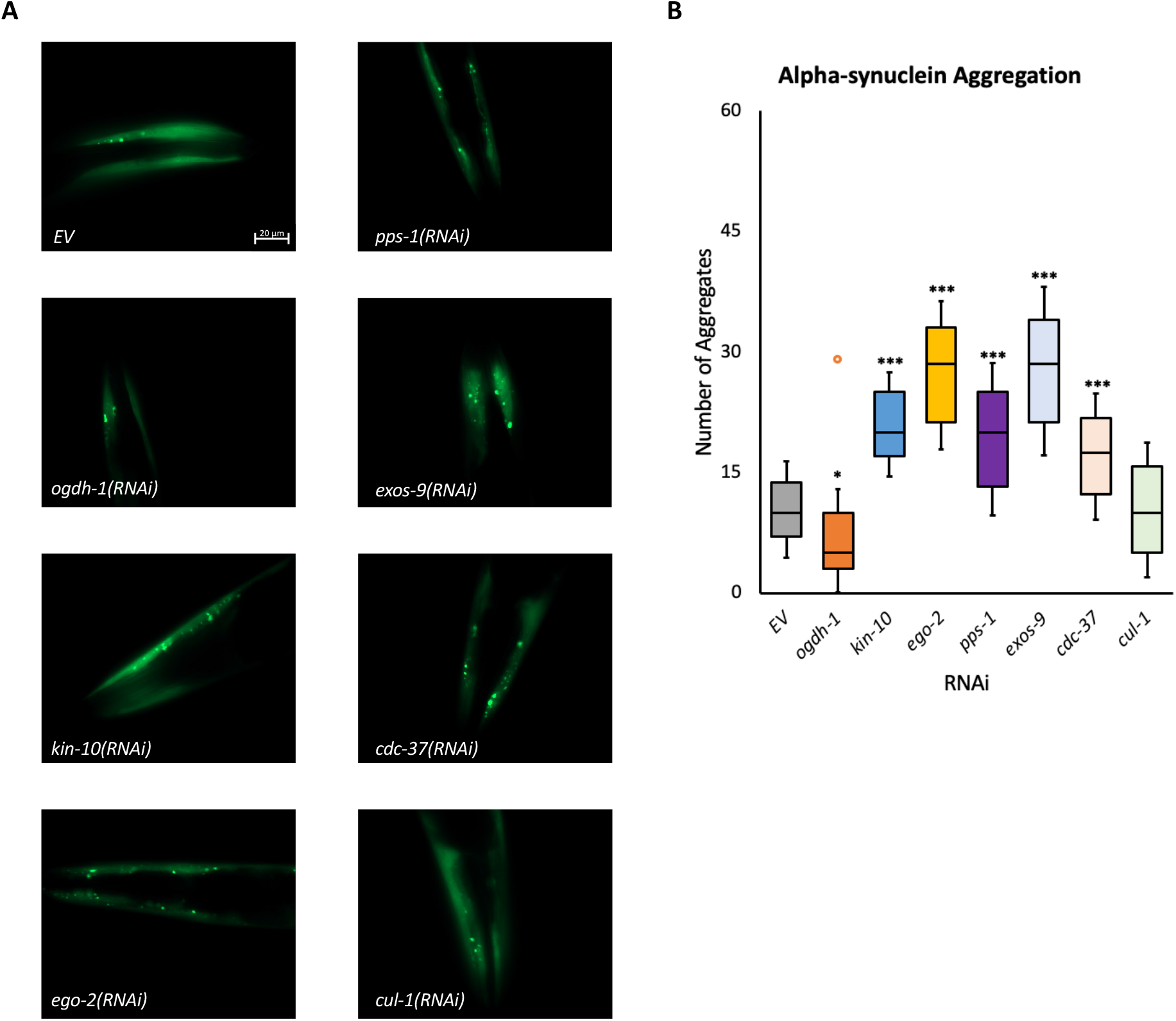
Genes in the mitochondrial health gene network limit proteostatic aggregation. **(A)** Representative fluorescence micrographs of a transgenic *C. elegans* strain containing an α-synuclein::YFP protein in worms reared on RNAi targeting the indicated gene. **(B)** Quantification of α-synuclein::YFP aggregates in young adult worms reared on the indicated RNAi. Statistical significance was calculated using one-way ANOVA analysis followed by a Dunnett’s post-hoc test. *p<0.05, **p<0.01, ***p<0.001

### Pathways related to human disease and biological processes related to cellular division are overrepresented amongst human orthologs

To extend the utility of our gene network, we looked for putative human orthologues. Since it is common for gene duplication events to have expanded gene families in humans compared to *C. elegans*, a combination of tools such as OrthoList, DIOPT, and Wormbase, in addition to manual curation were used for putative ortholog assignment. We attempted to strike a balance between thoroughness and not including an overabundance of genes with highly similar co-orthologs (e.g. histone proteins, certain HSPs, *etc*), so only the two human genes with the greatest mutual similarity were considered putative co-orthologs of *C. elegans* gene **(Supplemental Table 3)**. The resulting human network was comprised of 156 genes with 503 connections (*p-value*<1.00E-16) **(Figure 7A)**. A pull of 156 random genes from the human genome generates a network with only 37 connections (*p-value*=0.37) **(Figure 7B)**. Gene ontology enrichment analysis using the KEGG database via WebGestalt identified several categories that were overrepresented, including cell cycle, signaling pathways (e.g., Wnt, Hedgehog, Notch), and autophagy **(Figure 7C)**. Given that cancer cells were used for the identification of the genes tested for the initial seed set, the presence of these pathways was relatively unsurprising. However, this does not explain why so many of the genes identified disrupted mitochondrial function when mutated. It suggests that identified genes may play unappreciated roles in mitochondrial function as well, providing a tantalizing avenue for research. The presence of various diseases in the previous enrichment analysis led us to conduct an enrichment analysis focused solely on disease. For this bioinformatic analysis, we used the OMIM expanded database which contains extensive data on human genes and their phenotypes and can be filtered by subtype including disease (Amberger *et al*. 2015). Using the Enrichr analysis tool along with the Enrichr Appyter function, we generated a scatterplot of diseases that were enriched in our gene network **(Figure 7D)**. The diseases were clustered by genetic similarity, and we also obtained a list of the enriched diseases along with significance scores **(Supplemental Table 4)**. Even though the initial gene list was derived from cancer cell line mutations, the list of enriched diseases from our final network is comprised of a wide range of diseases such as lissencephaly, osteoporosis and coloboma. This further validates the notion that mitochondrial dysfunction can lead to a plethora of cellular malfunctions and further study of this network can potentially uncover novel mechanisms for mitochondrial homeostasis.

**Figure 7.**
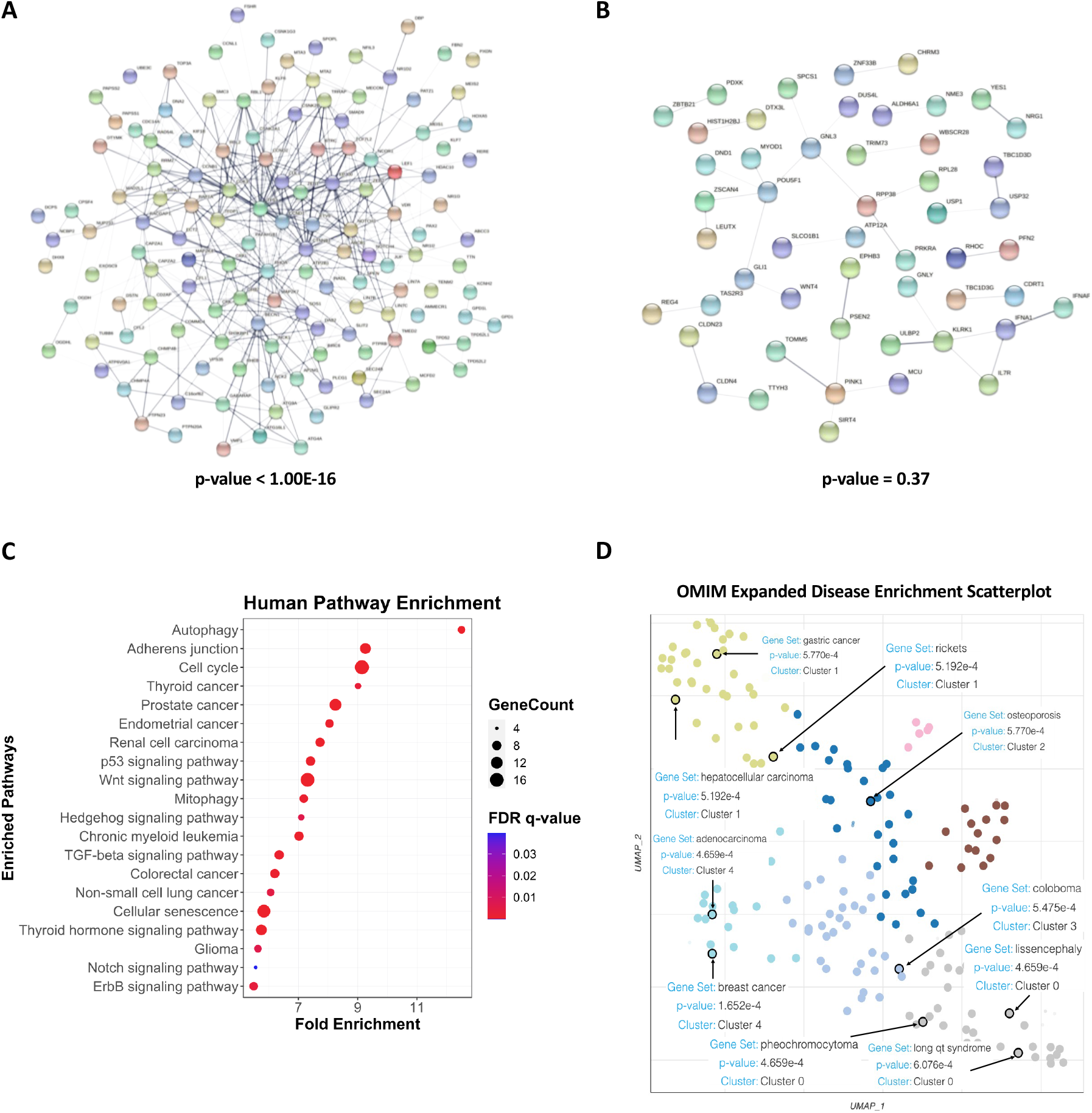
Human orthologs of the *C. elegans* mitochondrial health gene network illustrate increased co-association and are associated with disease. **(A)** A network of human genes comprised of 153 orthologs of the C. elegans mitochondrial health gene network significant number of connections. **(B)** Connectivity of one representative set (of 100 sets) of 153 randomly selected genes from the human genome, showing a total of ^~^40 connections. *p*-values **(B, C)** were calculated by the STRING database. **(C)** Bioinformatic analysis of gene function for the human genes shows strong enrichment for autophagy and mitophagy as well as cancer and cell division. Several pro-growth signaling pathways (such as p53, Wnt, Hedgehog, TGF-beta, Notch, and ErbB signaling pathways) are also enriched. **(D)** Analysis of human orthologs also shows enrichment for a variety of diseases.

## DISCUSS1ON

In this manuscript we identified a network of 139 genes that are important for mitochondrial maintenance. This provides the first indication of this role for many of these genes and, in a broader context, validates the idea of screening cancer cells to identify genes that are involved in processes like mitochondrial maintenance.

Of the 8 genes tested in the panel, *pps-1(RNAi)* showed the greatest difference from empty vector controls **(Table 2)**. This gene encodes a synthase that produces 3’-phosphoadenosine-5’-phosphosulfate (PAPS), a key intermediate required for sulfation in eukaryotes (van den Boom *et al*. 2012). Under normal circumstances, PPS-1 converts ATP into PAPS; this likely explains the ATP buildup in *pps-1(RNAi)* mutants. PPS-1 is a required gene in *C. elegans*, and is the sole orthologue of two genes in humans, PAPSS1 and PAPSS2, that are both required and play non-redundant roles (van den Boom *et al*. 2012). Mutations in PAPSS1 and PAPSS2 are associated with a broad range of phenotypes, including defects in formation of bone and cartilage formation and hormone biosynthesis (Gunal *et al*. 2019).

Although the function of PPS-1 is known, the connection between sulfation and mitochondria remains unclear, in part because of the variety of sulfated substrates. In an attempt to get a hint, we analyzed the genes connected to *pps-1, pop-1, gdph-2, sulp-3, sulp-5, cul-1*, and *dtmk-1*. In this case, this did not provide a clear answer either. One of the genes, *pop-1*, is involved in β-catenin binding, enzyme binding and transcription corepressor binding and is an orthologue for human LEF1 and TCF7L2. Interestingly, overactivation of SIRT1, which binds and activates LEF1, drives increased mitochondrial biogenesis and reduced mitochondrial turnover, leading to increased ATP and ROS (Song and Hwang 2018). SIRT1 is the most prominent sirtuin in mammals and is responsible for deacetylation of various proteins involved in a myriad of pathways including metabolic regulation, cell cycle, apoptosis, and inflammation (Yeung *et al*. 2004; Yi and Luo 2010; Lu *et al*. 2014; Tang 2016). This presents the tantalizing possibility that there may be a link between PAPSS and sirtuin function in mitochondrial maintenance.

Phenotypic assays of the genes in the panel also showed a significant increase in α-synuclein aggregates in a *C. elegans* NDD model. While it is not a faithful biomarker *per se*, misfolding and aggregation of α-synuclein in neural tissue is a hallmark of Parkinson’s disease (Ganguly *et al*. 2021). Mitochondrial dysfunction is known to be implicated in PD, there is still much that is unknown about what specific genes and pathways contribute to the progression of this disease (Malpartida *et al*. 2021). Recent findings have indicated that downregulation of mitochondrial S1RT3 may play a role in the mitochondrial dysfunction in PD and that α-synuclein may play a part in the decrease in S1RT3 expression through a decrease of phosphorylated AMPKα and phosphorylated CREB (Park *et al*. 2020). As shown above, the mitochondrial dysfunction present within the RNAi we tested is very similar to the mitochondrial dysfunction observed within the disruption of genes within the sirtuin pathways (SIRT1 and S1RT3). While there are no known association between our validation gene panel and the aforementioned sirtuins, it may indicate a possibility of links between them. Our gene network is proving to be a good reference for genes responsible not only for mitochondrial maintenance, but also for genes implicated in a variety of diseases.

In addition to measuring the phenotypic effects of knocking down the genes in our panel, we examined the network for gene connectivity and to determine whether any of the genes identified were plausible global regulators of mitochondrial maintenance. The most connected gene in our network was *rnr-2*, which encodes a subunit of ribonucleotide reductase, a cell cycle gene required for DNA synthesis orthologous to human RRM2. Human RRM2 is known to be implicated in cancer and has functions that range from the regulation of BRCA1 regulation to protection against ferroptosis (Rasmussen *et al*. 2016; Yang *et al*. 2020). While there is an extensive amount of literature regarding RRM2 and cancer, very little is known of the connection between RRM2 and mitochondrial maintenance. RRM2 is associated with a rare disease known as mitochondrial DNA depletion syndrome (MDS) and is linked to mitochondrial DNA (mtDNA) copy number in various models (Bornstein *et al*. 2008; Ylikallio *et al*. 2010; Villarroya *et al*. 2011). MDS is marked by malfunctions in mtDNA synthesis, reduced mitochondrial quantity, host clinical symptoms and high infant mortality rates, possibly due to depletion of nucleotide pools that RRM2 generates. Our network indicates that RRM2 is a hub for many of the genes in our network, potentially suggesting a larger role for RRM2 in mitochondrial maintenance than previously understood.

Overall, our findings provide insight into the genes and pathways that are needed for proper mitochondrial functions. Some of the network members were previously not know to have roles in mitochondrial homeostasis. Out of the 139 genes from our network, 101 of them have no known relationship to mitochondrial function. This further exemplifies the need for a better understanding and discovery of genes related to mitochondrial function and overall cellular maintenance. Knockdown of multiple pathway members from the test panel disrupted mitochondrial function (e.g. fission-fusion imbalance, changes in mitochondrial mass, NADH, ROS, etc.) with concomitant phenotypic consequences (changes in pumping and/or brood size), including exacerbation of protein proteostatic defects. The immediate value of this network is providing new fundamental knowledge about mitochondrial health. Additionally, however, they may also be useful for assessing physiological manifestations of mitochondria-related disruption in diseases that may not be intuitively connected to mitochondrial dysfunction, such as proteostatic disorders and cancer.

## ACKNOWLEDGEMENTS

Authors would like to thank Quinton L. Anderson (Rice University) for the help at the early stages of the project and Yizhi (Jane) Tao (Rice University) for providing some of the RNAi bacteria. NVK, a CPRIT scholar in Cancer Research, thanks the Cancer Prevention and Research Institute of Texas (CPRIT) for their generous support, CPRIT grant RR150044. This work was also supported by the National Institutes of Health (NIGMS R35GM129294) to NVK.

## SUPPLEMENTARY MATERIAL

### Supplemental Figures

**Supplemental Figure 1. Examples of screening assays outputs**. Representatives of assay hits or genes that didn’t pass a cutoff are shown. **(A)** In the precocious mitophagy activation assay (primary screen), genes where RNAi knockdown that exhibited a 1.5-fold increase in fluorescence compared to the empty vector control (indicating precocious activation of mitophagy) were selected as primary hits. **(B)** In the acute mitochondrial damage assay (secondary screen), genes where RNAi knockdown that exhibited a 1.5-fold increase in worm death compared to the empty vector control were selected as secondary hits.

**Supplemental Figure 2. Knockdown of majority of panel members results in abnormal mitochondrial membrane potential**. MitoTracker Red CMXRos was used to measure mitochondrial membrane potential in a RNAi knockdowns of panel members. Statistical significance was calculated using ANOVA with Dunnett’s *post hoc* test. * *p*<0.05, ** *p*<0.01, *** *p*<0.001

### Supplemental Tables

**Supplemental Table 1. List of 139 members of *C. elegans* mitochondrial health network**.

Seed and expanded set are marked by green and blue color, respectively. Known association with mitochondrial function is marked.

**Supplemental Table 2. List of 400 *C. elegans* genes with a significant connection to seed set**.

400 genes that are highly connected to the initial gene set were obtained using WormNetV3. Connections are considered significant connectivity *p*-value calculated by WormNetV3 were <0.05.

**Supplemental Table 3. List of human orthologs for 139 *C. elegans* genes of expended network**.

Orthologs were identified using OrthoList, DIOPT, and Wormbase and manually curated.

**Supplemental Table 4. List of enriched OMIM categories**.

OMIM disease enrichment shows a wide variety of diseases that are enriched in the mitochondrial health gene network. Statistical significance (*p*-values) and false discovery rate (*q* values) are shown.

