## Supplementary figures and images for "A Novel *Caenorhabditis elegans* Gene Network Uncovers Mechanisms of Mitochondrial Maintenance"

### Supplemental Figures

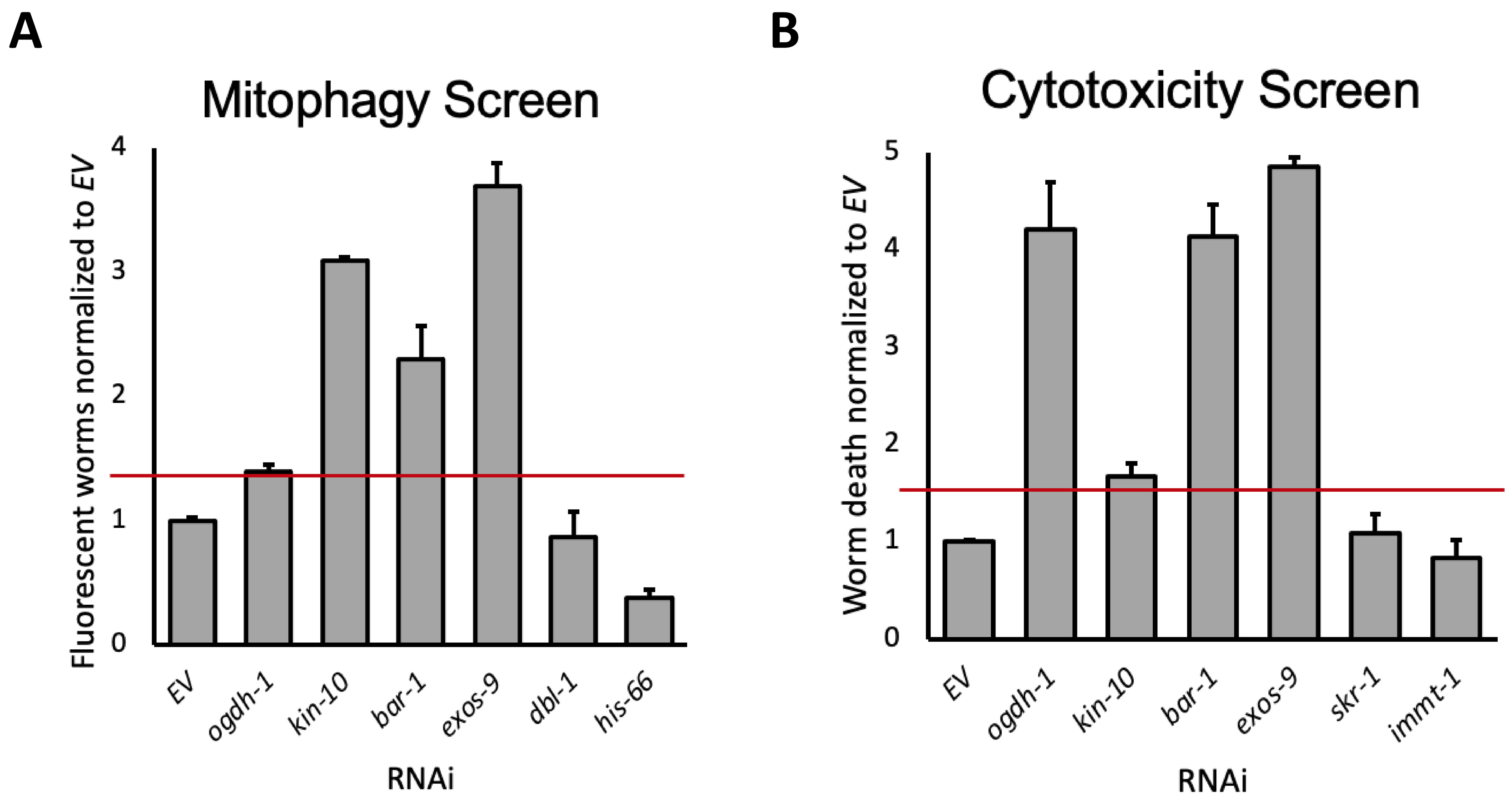


#### Supplementary Figure 1

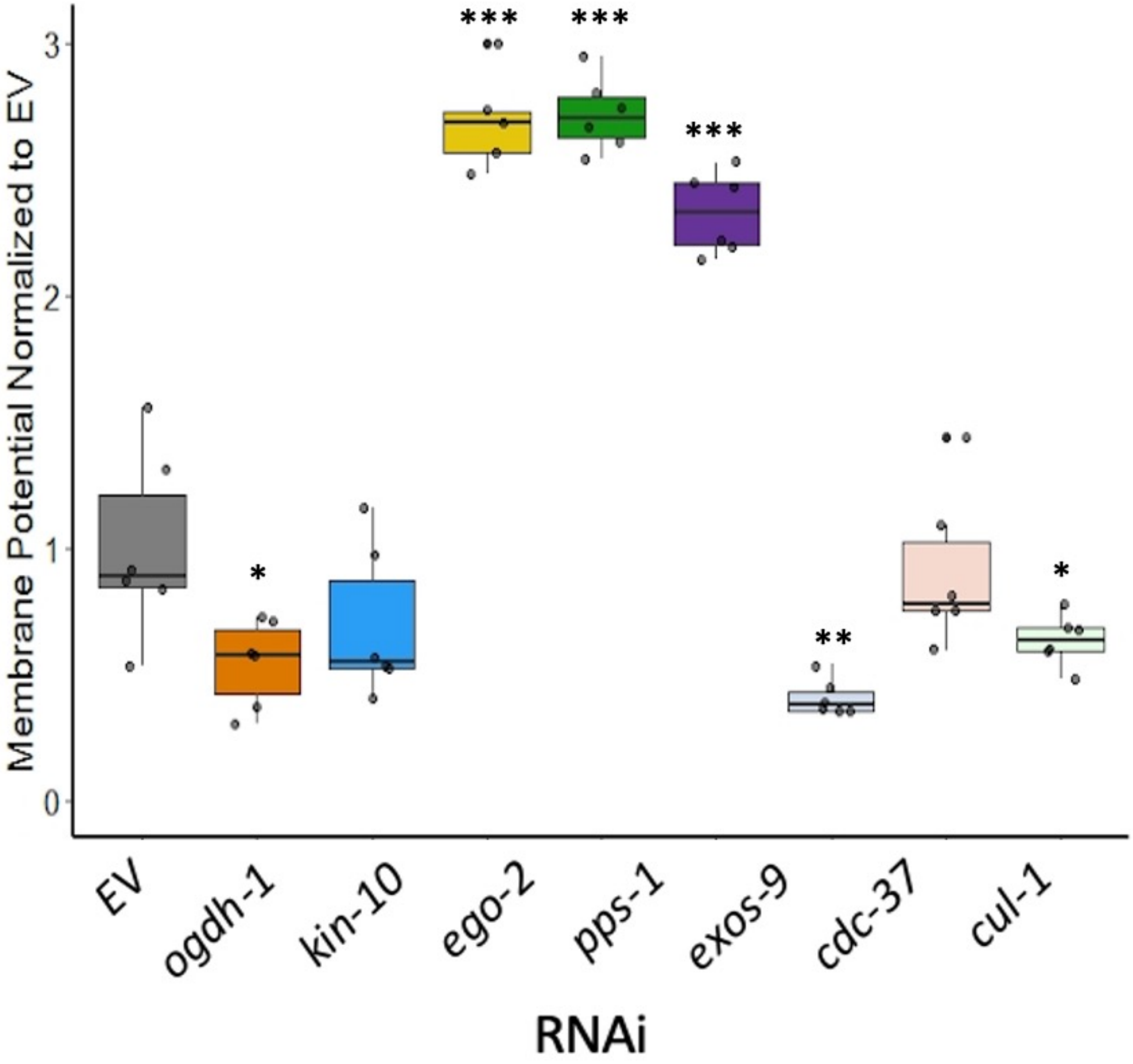


Supplemental Figure 2
